# Challenges and recommendations to improve installability and archival stability of omics computational tools

**DOI:** 10.1101/452532

**Authors:** Serghei Mangul, Thiago Mosqueiro, Richard J. Abdill, Dat Duong, Keith Mitchell, Varuni Sarwal, Brian Hill, Jaqueline Brito, Russell Jared Littman, Benjamin Statz, Angela Ka-Mei Lam, Gargi Dayama, Laura Grieneisen, Lana S. Martin, Jonathan Flint, Eleazar Eskin, Ran Blekhman

**Affiliations:** Department of Computer Science, University of California Los Angeles, 580 Portola Plaza, Los Angeles, CA 90095, USA; Institute for Quantitative and Computational Biosciences, University of California Los Angeles, 611 Charles E. Young Drive East, Los Angeles, CA 90095, USA; Department of Genetics, Cell Biology and Development, University of Minnesota, 321 Church St SE, Minneapolis, MN 55455, USA; Indian Institute of Technology Delhi, Hauz Khas, New Delhi, Delhi 110016, India; Institute of Mathematics and Computer Science, University of São Paulo, São Paulo, Brazil; Center for Neurobehavioral Genetics, Semel Institute for Neuroscience and Human Behavior, University of California Los Angeles, 760 Westwood Plaza, Los Angeles, CA 90095, USA; Department of Human Genetics, University of California Los Angeles, 695 Charles E. Young Drive South, Los Angeles, CA 90095, USA; Department of Ecology, Evolution, and Behavior, University of Minnesota, 100 Ecology Building, 1987 Upper Buford Cir, Falcon Heights, MN 55108, USA

## Abstract

Developing new software tools for analysis of large-scale biological data is a key component of advancing modern biomedical research. Scientific reproduction of published findings requires running computational tools on data generated by such studies, yet little attention is presently allocated to the installability and archival stability of computational software tools. Scientific journals require data and code sharing, but none currently require authors to guarantee the continuing functionality of newly published tools. We have estimated the archival stability of computational biology software tools by performing an empirical analysis of the internet presence for 36,702 omics software resources published from 2005 to 2017. We found that almost 28% of all resources are currently not accessible through URLs published in the paper they first appeared in. Among the 98 software tools selected for our installability test, 51% were deemed “easy to install,” and 28% of the tools failed to be installed at all due to problems in the implementation. Moreover, for papers introducing new software, we found that the number of citations significantly increased when authors provided an easy installation process. We propose for incorporation into journal policy several practical solutions for increasing the widespread installability and archival stability of published bioinformatics software.

## Introduction

During the past decade, the rapid advancement of genomics and sequencing technologies has inspired a large and diverse collection of new algorithms in computational biology ^1, 2^. In the last 15 years, the amount of available genomic sequencing data has doubled every few months ^3, 4^. Life-science and biomedical researchers are leveraging computational tools to analyze this unprecedented volume of genomic data ^3, 4^, which has been critical in solving complex biological problems and subsequently laying the essential groundwork for the development of novel clinical translations ^5^. The exponential growth of genomic data has reshaped the landscape of contemporary biology, making computational tools a key driver of scientific research ^6, 7^.

As computational and data-enabled research become increasingly popular in biology, novel challenges arise, accompanied by standards that attempt to remedy them. One such challenge is computational reproducibility—the ability to reproduce published findings by running the same computational tool on the data generated by the study ^8–10^. While several journals have introduced requirements for the sharing of data and code, there are currently no effective requirements to promote installability and long-term archival stability of software tools, creating situations in which researchers share source code that either doesn’t run or disappears altogether. These issues can limit the applicability of the developed software tools and impair the community’s ability to reproduce results generated by software tools in the original publication.

The synergy between computational and wet-lab researchers is especially productive when software developers distribute their tools as packages that are easy to use and install ^11^. Though many new tools are released each year, comparatively few incorporate adequate documentation, presentation and distribution, resulting in a frustrating situation in which existing tools address every problem except how to run them ^12^.

Widespread support for software installability promises to have a major impact on the scientific community ^13^, and practical solutions have been proposed to guide the development of scientific software ^14, 16–17^. While the scale of this issue in computational biology has yet to be estimated, the bioinformatics community warns that poorly maintained or improperly implemented tools will ultimately hinder progress in data-driven fields like genomics and systems biology ^3, 7, 18^.

### Challenges to effective software development and distribution in academia

Successfully implementing and distributing software for scientific analysis involves numerous unique challenges that have been previously outlined by other scholars ^11, 15, 16, 19^. In particular, fundamental differences between software development workflows in academia and in industry challenge the installability and archival stability of novel tools developed by academics. These differences can be broken down into three broad categories:

- Software written by researchers tends to be written with the idea that users will be knowledgeable about the code and appropriate environment and dependencies. This sometimes results in tools that are difficult to install, with instructions and command-line options that are unclear and confusing, but are also critical for the tool’s function.
- Academic journals are a primary source for information and documentation of noncommercial scientific software, even though the static nature of publications means this vital information quickly falls out of date.
- Incentives in academia heavily favor the publication of new software, not the maintenance of existing tools.

First, software developers in industrial settings receive considerably more resources for developing user-friendly tools than their counterparts in academic settings ^20^. Commercial software is developed by large teams of software engineers that include specialized user experience (UX) developers. In academic settings, software is developed by smaller groups of researchers who may lack formal training in software engineering, particularly UX and cross-platform design. Many computational tools lack a user-friendly interface to facilitate the installation or execution process ^12^. Developing an easy-to-use installation interface is further complicated when the software relies on third-party tools that need to be installed in advance, called “dependencies.” Installing dependencies is an especially complicated process for researchers with limited computational knowledge. Well-defined UX standards for software development could help software developers in computational biology promote widespread implementation and use of their newly developed computational tools.

Second, companies efficiently distribute industry-produced software using dedicated company units or contractors—services that universities and scientific funding agencies do not typically provide for academic-developed software. The computational biology community has adopted by default a pragmatic, short-term framework for disseminating software development,^21^ which generally consists of publishing a paper describing the software tool in a peer-reviewed journal. So-called “methods papers” are dedicated to explaining the rationale behind the novel computational tool and demonstrating its efficacy with sample datasets. Supplemental materials such as detailed instructions, tutorials, dependencies, and source code are made available on the internet and included in the published paper as a URL, but generally exist in a location out of the journal’s direct control. The quality, format, and long-term availability of supplemental materials varies among software developers and is subject to less scrutiny in the peer-review process compared to the published paper itself. This approach limits the installability of software tools for use in research and hinders the community’s ability to evaluate the tools themselves ^22^.

Third, the academic structures of funding, hiring, and promotion offer little reward for continuous, long-term development and maintenance of tools and databases ^23^, and software developers can lose funding for even the most widely used tools. Loss of external funding can slow and even discontinue software development, potentially impacting the research productivity of studies that depend on these tools ^24^. Interrupted development also hinders the ability to reproduce results from published studies that use discontinued tools. In general, industry-developed software is supported by teams of software engineers dedicated to developing and implementing updates for as long as the software is considered valuable. Many software developers in academia do not have access to mechanisms that could ensure a similar level of maintenance and stability.

We combined two approaches to determine the effects of these challenges on the proportion of bioinformatics tools that could be considered user-friendly. First, we investigated tens of thousands of URLs corresponding to bioinformatics tools and resources to determine whether they are archivally stable—whether users can even reach the websites described in the papers evaluated. Next, we investigated the number of tools that provided an easy-to-use installation interface to download and install the software and any required dependencies.

### Archival stability of published computational tools and resources

The World Wide Web provides a platform of unprecedented scope for data and software archival stability, yet long-term preservation of online resources remains a largely unsolved problem ^25^. Published software tools are made accessible through the Uniform Resource Locator (URL), which is typically provided in the abstract or main text of the paper and is often assumed to be a practically permanent locator. However, a URL may become inactive due to removal or reconfiguration of web content. The “death of URLs” ^26^ has been described for decades in various terms, including “link rot” ^27^ and “lost Internet References.” ^28^ At the onset, the World Wide Web promised the virtually infinite availability of digital resources; in practice, many are lost. For example, many tools in computational biology are hosted on academic web pages that become inactive with time, sometimes only months after their initial publication. These software packages are typically developed by small groups of graduate students or postdoctoral scholars who, considering the temporary nature of such positions, cannot maintain such websites and software for longer periods of time.

Multiple studies have identified the deterioration of long-term archival stability of published software tools ^18, 26, 28–31^. In order to begin assessing the magnitude of these issues in computational biology, we comprehensively evaluated the archival stability of computational biology tools used in 51,236 biomedical papers published across 10 relevant peer-reviewed journals over a span of 18 years, from 2000 to 2017 (Table S1). Out of the 51,236 examined papers, 13.6% contained at least one URL in their abstracts, and another 38.3% contained URLs in the body of the paper. To increase the likelihood that the identified URL corresponds to a software tool or database, we inspected 10 neighboring words for specific keywords commonly used, including “pipeline”, “code”, “software”, “available”, “publicly”, and others (See Methods Section). Complete details on our methodology for extracting the URLs, including all parameters and thresholds, are provided in the Supplementary Methods.

We used a web mining approach to test 36,702 published URLs that our survey identified. We categorized unreachable URLs into two groups: unreachable due to connection timeout and unreachable due to error (“broken” links, i.e., 404 HTTP status). We separately categorized accessible URLs that returned immediately and those that used redirection—that is, URLs to which servers responded by pointing the user to a new URL that then connects successfully. We found that 26.7% of evaluated URLs are successfully redirected to new URLs. (Some URLs were redirected to pages that subsequently returned an error; these were not considered successful.) Of all identified URLs, 11.9% were unreachable because of connection timeouts, and 15.9% were “broken.” To prevent erroneous classification caused by configuration of our automated tests, we manually verified more than 900 URLs reported as “timeouts,” or requests that did not receive a response within an acceptable amount of time (Figure S1).

Next, we grouped the URLs by the year in which the computational biology tool was first referenced in a publication. As expected, the time since publication is a key predictor of URL archival stability (Fisher exact test, p-value <10^−15^). 41.9% of the software referenced before 2012 (n=15,439) is unavailable, whereas only 17.5% of the recent software referenced in 2012 and later (n=21,263) is unavailable (Figure 1a). After 2013, we observe a drop in the absolute number of archivally unstable resources (Figure 1b). Despite the strong decline in the percentage of missing resources over time, there are still 200 resources published every year with links that were broken by the time we tested them. The data and scripts for reproducing the plots in Figure 1 are available at https://github.com/smangul1/good.software.

**Figure 1.**
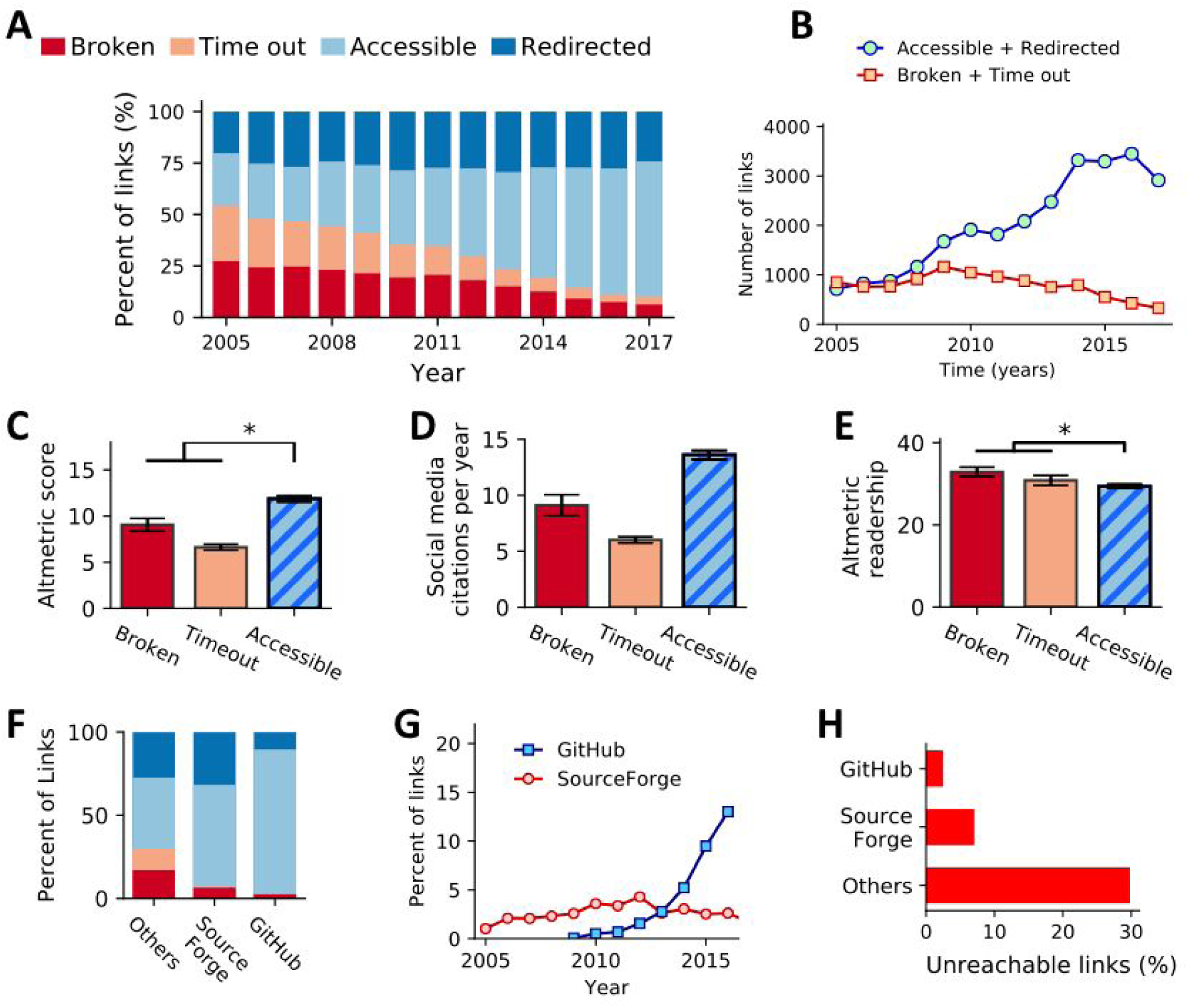
Archival stability of 36,702 published URLs across 10 systems and computational biology journals over the span of 13 years. An asterisk (*) denotes categories that have a difference that is statistically significant. Error bars, where present, indicate standard error of the mean (SEM). **(a)** Archival stability status of all links evaluated from papers published between 2005 and 2017. Percentages of each category (y-axis) are reported over a 13-year span (x-axis). **(b)** A line graph comparing the overall numbers (y-axis) of functional (green circles) and non-functional (orange squares) links observed in papers published over time (x-axis). **(c)** A bar chart showing the mean Altmetric “attention score” (y-axis) for papers, separated by the status of the URL (x-axis) observed in that paper. **(d)** A bar chart showing the mean number of mentions of papers in social media (blog posts, Twitter feeds, etc.) according to Altmetric, divided by the age of the paper in years (y-axis). Papers are separated by the status of the URL (x-axis) found in the paper. **(e)** A bar chart illustrating the mean Altmetric readership count per year of papers (y-axis) containing URLs in each of the categories (x-axis). **(f)** The proportion of unreachable links (due to connection timeout or due to error) stored on web services designed to host source code (e.g., GitHub and SourceForge) and ‘Other’ web services. **(g)** A line plot illustrating the proportion (y-axis) of the total links observed in each year (x-axis) that point to GitHub or SourceForge**. (h)** A bar chart illustrating the proportion of links hosted on GitHub or SourceForce (vertical axis) that are no longer functional (horizontal axis), compared to links hosted elsewhere.

Prior research demonstrates that the availability of published bioinformatics resources has a significant impact on citation counts ^30^. In addition to those generally accepted measures of scientific impact, we assessed the effect of software availability on complementary metrics of impact, such as measures of social media mentions, media coverage, and public attention (Figure 1c–e). We found that papers with accessible links exhibit increased engagement by readers in social media, reflected in a significantly higher number of citations in social media platforms per year (Figure 1d; Kruskal–Wallis *p-*value = 1.75×10^−161^, Dunn’s test p=9.66×10^−103^ for accessible vs. broken, p=3.37×10^−76^ for accessible vs. timeout, adjusted for multiple tests using the Benjamini–Hochberg procedure) and an increased Altmetrics score ^32^ when compared to papers with “broken” and “timeout” links (Figure 1c; Kruskal–Wallis, *p-*value = 1.66×10^−25^, Dunn’s test p=2.47×10^−17^ for accessible vs. broken, p=4.16×10^−14^ for accessible vs. timeout). While the difference is small, we found the readership of papers with accessible links differed significantly from papers with links that are classified as broken or timeouts—surprisingly, the median reader count per year (according to Altmetric) was lower for papers with accessible links (Figure 1e; Dunn’s test p=1.17×10^−6^ for accessible vs. broken, p=8.42×10^−15^ for accessible vs. timeout).

In addition, we tested the impact of using websites designed to host source code, such as GitHub and SourceForge, on the archival stability of bioinformatics software. These websites have been used by the bioinformatics community since 2001, and the proportion of software tools hosted on these sites has grown substantially, from 1.6% in 2012 to 13% in 2016 (Figure 1g). We find that URLs pointing to these websites have a high rate of archival stability: 97.6% of the links to GitHub and 93.0% of the links to Sourceforge are accessible, while only 70.3% of links hosted elsewhere are accessible (Figure 1h), a significant difference (Fisher-exact test GitHub vs. Others, p=2.70×10^−106^; SourceForge vs. Others p=1.31×10^−5^).

Our results suggest that the computational biology community would benefit from such approaches, which effectively guarantee permanent access to published scientific URLs. Specifically, several key principles emerge that promise to positively impact the availability of published bioinformatics resources, including the number of citations and social media references. In addition, bioinformatics tools and resources stored on web services designed to host source code have a significantly higher chance of remaining accessible.

### Tool installability

We developed a computational framework capable of systematically verifying the archival stability and installability of published software tools. We applied this framework to 98 randomly selected tools across various domains of computational biology (**Method Section**). Those tools were selected independently from the 36,702 URLs used above (‘Archival stability of published computational tools and resources’). We engaged undergraduate and graduate students to run the installation test using a standardized protocol (Figure S2); we recorded the time required to install the tools and other important features, allowing up to two hours per software package. In total, 71 hours of installation time was required in attempts to install 98 tools. We categorized a tool as “easy to install” if it could be installed in 15 minutes or less, “complex” if it required more than 15 minutes but was successfully installed before the two hour limit, and “not installed” if the tool could not be successfully installed within two hours (Table S2 and Figure 2).

**Figure 2.**
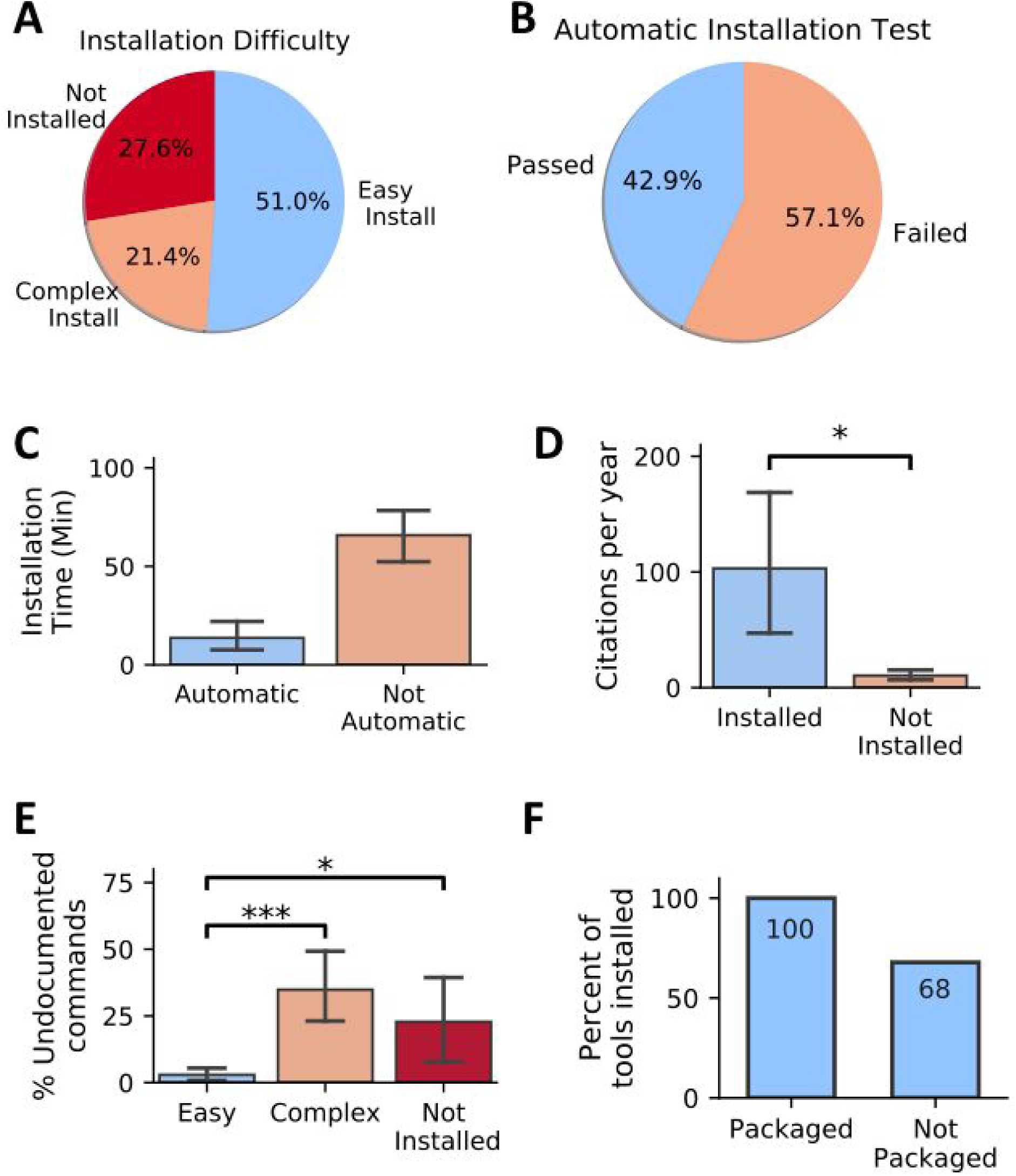
Installability of 98 randomly selected published software tools across 22 life science journals over a span of 15 years. Error bars, where present, indicate SEM. **(a)** Pie chart showing the percentage of tools with various levels of installability. **(b)** A pie chart showing the proportion of evaluated tools that required no deviation from the documented installation procedure. **(c)** Tools that require no manual intervention (pass automatic installation test) exhibit decreased installation time. **(d)** Tools installed exhibit increased citation per year compared with tools which were not installed (Kruskal–Wallis, *p-*value = 0.035). **(e)** Tools which are easy to install include a decreased portion of undocumented commands (Not Installed vs. Easy Install: Mann–Whitney *U* test, p-value=0.01, Easy Install vs. Complex Install: Mann–Whitney *U* test, p-value=8.3×10^−8^). **(f)** Tools available in well-maintained package managers such as Bioconda were always installable, while tools not shipped via package managers were prone to problems in 32% of the studied cases.

The most stringent evaluation was the “automatic installation test,” in which the tester is required to strictly follow the instructions provided in the manual of the software tool (Methods; Figure 2a)—we determined that 57.1% of the selected tools failed this test. The vast majority (39 out of 42) of the tools that passed this test finished in fewer than 15 minutes and were classified as “easy to install” (Table S2). For the tools failing the test, we performed manual intervention during which the tester was allowed to install missing dependencies and modify code to resolve installation errors. On average, it took an additional 70 minutes to install tools failing the “automatic installation test” (Mann–Whitney *U* test, p-value=4.7×10^−9^; Figure 2c). Manual intervention was unsuccessful for 66% of the tools that initially failed the automatic installation test. Failed manual installation was due to numerous issues, including hard-coded parameters, invalid folder paths or header files, and usage of unavailable software dependencies.

Next, we assessed the effect of the ease of installation on the popularity of tools in the computational biology community by investigating the number of citations for the paper describing the software tools. We find that tools which we were able to install had significantly more citations compared to tools which we were not able to successfully install within two hours (Figure 2d; Mann–Whitney *U* test, p-value=0.032). These results suggest, perhaps not surprisingly, that tools which are easier to install are more likely to be adopted by the community.

In addition, we aimed to see whether the accuracy of a tool’s installation instructions affects its installation time. Considering the proportion of commands that are undocumented (estimated as a ratio between the executed commands and commands in the manual), we find that tools with easier installation have a significantly lower percentage of undocumented commands (Figure 2e; Mann–Whitney *U* test, p-value=0.04). Considering a significant increase of installation time and a low rate of success for tools failing automatic installation test, we argue that reliance on manual intervention to successfully install and run computational biology tools is an unsustainable practice. Software developers would benefit from ensuring a simple installation process and providing adequate installation instructions.

The vast majority of surveyed tools fail to provide one-line solutions for installation, instead providing step-by-step instructions. On average, eight commands were required to install surveyed tools, while only 3.9 commands were provided in the manual. Among the surveyed software tools, 23 tools provide one-line installation solutions that worked successfully, of which nine were available via the Bioconda package manager ^33^ (Table S2). A package manager is a system that automates the installation, upgrade, and configuration of a collection of software tools in a consistent manner. Tools with single-command installations require on average 6 minutes of installation time, which is significantly faster when compared to tools which require multi-command installation (Kruskal–Wallis, p-value=4.7×10^−6^) (Figure S3). Tools available in well-maintained package managers (e.g., Bioconda) were always installable, while tools not shipped via package managers failed to install in 32% of the studied cases (Figure 2f).

### Box 1. Principles to increase installability and archival stability of omics computational tools and resources.

The results from our study point to several specific opportunities for establishing an effective software development and distribution practice. Here we present five principles to increase the installability and archival stability of omics computational tools and resources. The majority of surveyed software tools and resources address only a portion of these principles.

1. **Host software and resources on archivally stable services** Selecting the appropriate service to host your software and resources is critical. A simple solution is to use web services designed to host source code (e.g. GitHub ^34, 35^ or Sourceforge). In our study, we have determined that more than 96% of software tools and resources stored at GitHub or Sourceforge are accessible, and tools hosted on these services remain stable for longer periods of time (Table S3). Ideally, the repositories storing code should also be more permanently archived using a service such as Zenodo (https://zenodo.org), which is designed to provide long-term stability for scientific data.
2. **Provide easy-to-use installation interface** Use sustainable and comprehensive software distribution. One example of a sustainable package manager is Bioconda^33^, which is language-agnostic and available on Linux and Mac operating systems. Bioconda, technically a “channel” within the broader Conda project, is one the most popular package managers in bioinformatics, currently covering 2900 software tools that are continuously maintained, updated, and extended by a growing global community ^33^. Bioconda provides a one-line solution for downloading and installing a tool.
3. **Take care of all the dependencies the tool needs** Even the most widely used tools rely on dependencies. To facilitate simple installation, provide an easy-to-use interface to download and install all required dependencies. Ideally, all necessary installation instructions should be included in a single script, especially when the number of installation commands is large. Package managers can potentially make this problem easier to solve. Bioconda also automatically generates containers for each Bioconda “recipe” ^36^, which provides all files and information needed to install a package. Other implementations of containerized software (e.g. Docker – https://www.docker.com/ and Singularity – https://www.sylabs.io/docs/) also usually have all dependencies preinstalled. Often, language-specific solutions are also available (e.g Bioconductor^37^ and the Comprehensive R Archive Network (CRAN)). One drawback of Bioconda is that the existing tools in portable package managers are manually updated by the team or community, often delaying such updates. For example, as of August 10, 2018, R 3.5 was unavailable under Bioconda despite being released almost four months prior. If possible, one should design an installation script combining the commands for installing dependencies and developed software tools into a single script. Ideally, these dependencies should be installed in a user-configurable directory, as with Python “virtual environments,” which can help avoid conflicts with existing software on the system.
4. **Provide an example dataset** Provide an example dataset inside the software package, with a description of the expected results. Similar to unit- and integration-testing practices in software engineering, example datasets allow the user to verify that the tool was successfully installed and works properly before running the tool on experimental data. A tool may be installed with no errors, yet it may still fail to successfully run on the input data. Only 68% of examined tools provide an example dataset (Table S2).
5. **Provide a ‘Quick Start’ guide** Allow the user to verify the installation and performance of the tool. Providing a ‘Quick Start’ guide is the best way for the user to validate that the tools are installed and working properly. The guide should provide the commands needed to download, install, and run the software tool on the example dataset. An example of a ‘Quick Start’ guide is provided in Supplemental Note 2. In addition to the ‘Quick Start’, a detailed manual must be provided with information on options, advanced features and configuration. Best practices for creating bioinformatics software documentation are discussed elsewhere ^38^.
6. **Choose an adequate name** Choose a software name that best reflects the developed tool or resource. Today’s “age of Google” places new demands on the function of tool names, which should be memorable and unique, yet easily searchable. In addition, there are no regulations on tool names. For example, there are at least six tools named “Prism,” making it challenging to find the right tool (Supplementary Note 3). Scout the web to check the uniqueness of a name before publishing a new tool.
7. **Assume no root privileges** Tools are often installed on high-performance computing clusters where users do not have administrative (root/superuser) privileges to install software into system directories. When developing instructions for installation of the proposed software tool, avoid commands that require root access. Examples of such commands include those that use package managers that require root/superuser privileges, such as “apt-get install” or “yum install.”
8. **Make platform-agnostic decisions when possible** Create tools that will work on as many systems as possible—specification of various versions of UNIX-like systems may limit the installability of software. Design your software to minimize reliance on OS-specific functionality to make it easier for users to use your tools in diverse environments. Platform-specific installation commands (e.g., Homebrew ^39^) should also be avoided.

## Discussion

Our study assesses a critical issue in computational biology that is characterized by lack of standards regarding installability and long-term archival stability of omics computational tools and resources. Despite recent requirements on the behalf of journals to impose data and code sharing on published authors’ work, 27.8% of 36,702 omics software resources examined in this study are not currently accessible via the original published URLs. Among the 98 software packages selected for our installability test, 49.0% of omics tools failed our “easy-to-install” test. In addition, 27.6% of surveyed tools could not be installed due to severe problems in the implementation process. One-quarter of examined tools are easy to install and use; in these cases, we identify a set of good practices for software development and dissemination.

Reviewers assessing the papers that present new software tools could begin addressing this problem with the adoption of a rigorous, standardized approach during the peer review process. Feasible solutions for improving the installability and archival stability of peer-reviewed software tools include requirements for providing installation scripts, test data, and functions that allow automatic checks for the plausibility of installing and running the tool. For example, “forking” is a simple procedure that ensures the version of cited code within an article may persist beyond initial publication ^40^. Academic journals recently took a major step toward improving archival stability by permanently forking published software on GitHub (e.g., ^41^).

The current workflow of computational biology software development in academia encourages researchers to develop and publish new tools, but this process does not incentivize long-term maintenance of existing tools. Results from this study provide a strong argument for the development of standardized approaches capable of verifying and archiving software. Further, our results suggest that funding agencies should emphasize support for maintenance of existing tools and databases.

Manual interventions and long installation times are unappealing to many users, especially to those with limited computational skills. Many life science and medical researchers lack formal computational training and may be unable to perform manual interventions (e.g., installing dependencies or editing computer code during installation). Users could leverage advanced knowledge of the time and computational skills required to properly install a software package. We propose a prototype of a badge server that runs an automated installation test, thus introducing to the peer review process explicit assessment of a tool’s installability. This badge server would be particularly useful in computational biology, an interdisciplinary field comprised of reviewers who often lack the skills and time to verify the installability of software tools. Many benchmarking studies already routinely report relative ease of installation and use of new tools as components of their performance metrics^42^.

## Acknowledgments

SM acknowledges support from a QCB Collaboratory Postdoctoral Fellowship, and the QCB Collaboratory community directed by Matteo Pellegrini. S.M. and E.E. are supported by National Science Foundation grants 0513612, 0731455, 0729049, 0916676, 1065276, 1302448, 1320589, 1331176, and 1815624, and National Institutes of Health grants K25-HL080079, U01-DA024417, P01-HL30568, P01-HL28481, R01-GM083198, R01-ES021801, R01-MH101782, and R01-ES022282. R.B. is grateful for support from the National Institutes of General Medicine (R35-GM128716) and a McKnight Land-Grant Professorship from the University of Minnesota. We thank John Didion (https://twitter.com/jdidion) for an interesting discussion over Twitter about the issue of software installability.

**Glossary**
installability: The tool is considered usable if (a) the tool and its corresponding dependencies can be installed on Linux/UNIX-based operating systems, and if (b) the tool can produce expected results from the input data with no errors.
Automated installation test: This test of software installation ease is performed by the biomedical researcher, using only installation commands provided in the manual in the recommended order. No extra commands are allowed. A tool passes the automated installation test if the user can successfully install the package following only the commands from the manual.
Package manager: is a collection of software tools that automate the installation of a tool’s core package and updates in a consistent manner. Package managers also help solve the ‘dependencies problem’ by automatically installing required third-party software packages. Bioconda is one of the most popular package managers for omics computational tools. A growing global community of Bioconda users continuously maintain, update, and extend more than 2900 software tools.

## Methods

### Protocol to check the archival stability of published software tools

We downloaded open access papers via PubMed from 10 systems and computational biology journals from the NCBI FTP server (ftp://ftp.ncbi.nlm.nih.gov/pub/pmc/). We included the following journals: *Nature Biotechnology, Genome Medicine, Nature Methods, Genome Biology, BMC Systems Biology, Bioinformatics, PLoS Computational Biology, BMC Bioinformatics, BMC Genomics*, and *Nucleic Acids Research*.

Papers were downloaded in XML format, which contains name-tags for field extraction. (Raw data from PubMed is available at https://doi.org/10.6084/m9.figshare.7641083) Specifically, we focused on three tags: <abstract>, <body>, and <text-link>. Each paper’s abstract is enclosed inside the <abstract> tag (Figure S1). The <body> tag contains the key contents like introduction, methods, results, and discussion. <ext-link> tags contain internet addresses for external sources (e.g., supplementary data and directions for downloading data sources and software packages). We have prepared a folder containing a small set of papers in XML format for testing purposes, available at https://github.com/smangul1/good.software/blob/master/download.parse.data/Nat_Methods.tar.gz?raw=true.

We deployed a heuristic approach to limit links to software produced by each paper’s authors. We assumed that these links are in <ext-link> tags whose neighbor words contain one of the following keywords: “here”, “pipeline”, “code”, “software”, “available”, “publicly”, “tool”, “method”, “algorithm”, “download”, “application”, “apply”, “package”, and “library”. We searched for these words in a neighborhood that extended 75 characters from both the start and end of each <ext-link> tag.

For each extracted link, we initially used the HTTPError class of the Python library urllib2 to get the HTTP status. Status number 400 and above indicate broken links; for example, the well-known 404 code indicates “Page Not Found.” Some URLs point at servers that did not respond at all. Since the threshold for the allotted time to wait for a response may bias the results, we manually verified 931 URLs reported with the timeout error code (Figure S1).

Multiple attempts were made to validate each extracted URL: First, an HTTP request was sent to each URL; if that was not successful, an FTP request was sent, to avoid marking URLs as “broken” if they used this older method of transferring files instead. HTTP requests that received “redirect” responses (status codes 300–399) were followed to the endpoint specified by the redirection (or redirections), to determine the final destination of the request. If the request ultimately completed successfully, the initial redirect code was recorded, and that link appears in our data as a redirection. However, some requests eventually resulted in errors—for example, if a server rewrites a received URL according to a formula, but the rewritten URL points to a file that doesn’t exist. Redirections that eventually resulted in an error were recorded with that error code instead. There is only one exception to this classification: If a server responded with a redirection status, but the redirection pointed at a URL that *only* changes the URL’s protocol from “http” to “https,” we classified this as a “success” rather than a “redirection.” Our protocol to check the archival stability of published software tools is available at https://github.com/smangul1/good.software. Parsed HTTP information for each links is available at https://doi.org/10.6084/m9.figshare.7738901.

### Protocol to check the installability of published software tools

To standardize the operating system environment for each tool installation, we used a CentOS 7 (v1710.01) Vagrant virtual machine. CentOS is an open-source operating system that is widely used in research computing. To prevent dependency mismatches caused by previously installed packages, we installed each tool in a new Vagrant virtual machine. Our virtual machine was provisioned with several commonly used software tools using the YUM package manager, to accommodate low-level dependencies that many developers would assume were already installed: epel-release, java (version 1.8.0), wget, vim, unzip, gcc and gcc-devel, python, and R. Users seeking to replicate this environment can use the Vagrant provisioning script found here: https://github.com/smangul1/good.software/blob/master/toolInstall/Vagrantfile

We present a summary of our protocol in Figure S3. Tools were classified into three categories: (1) easy to install, where installation took less than 15 minutes; (2) hard to install, where installation took between 15 minutes and two hours; and (3) not installed, meaning installation took longer than two hours or could not be completed. We tested a total of 98 tools across various categories and fields as described below. Information on the tools tested and the results of the test are available in **Table S2** and are shared at https://doi.org/10.6084/m9.figshare.7738949.

### Tools for microbiome profiling

The installability of 10 common tools for microbiome analysis was tested. To develop a list of popular tools, two co-authors independently made lists of 30 tools currently used for microbiome data processing, based on a literature survey, and identified those present on both lists. Microbiome tools can vary in their specificity of use; we limited the final tool list to five tools that process raw sequences into a final OTU table, and five tools capable of broad downstream analysis functions.

### Tools for read alignment

We tested the installability of 10 tools for read alignment. We randomly selected a total of 20 tools—10 tools from a recent survey ^43^ and 10 tools from PubMed (https://www.ncbi.nlm.nih.gov/pubmed/). The full list of extracted URLs is available at https://github.com/smangul1/good.software. To confirm that the installation process indeed worked, we used reads generated from the complete genome of Enterobacteria phage lambda (NC_001416.1).

### Tools for variant calling tools

We tested the installability of seven randomly sampled tools designed for variant calling ^44^. We confirmed successful software installation when the core functionality of each package could be executed with an example dataset. Only one of the tools was not packaged with an example dataset, in which case we randomly chose an open example dataset. We discarded from our study the tools for which papers could not be located.

### Tools for structural variants tools

We examined the installability of 52 common tools used for the structural variant (SV) calling from whole genome sequencing (WGS) data. First, we compiled a list of tools that use read alignment, where reads aligned to the locations are inconsistent with the expected insert size of the library or expected read depth at a specific locus. We randomly selected 50 tools out of 70 programs designed to detect SVs from WGS data and published after 2011. We confirmed the successful installation of each software package by executing its core functionality with an example dataset.

### Additional omics tools

Lastly, we randomly selected 20 published tools based on the URL present in the abstract or the body of the publications available in PubMed (https://www.ncbi.nlm.nih.gov/pubmed/). The full list of extracted URLs is available at https://github.com/smangul1/good.software.

### Statistical analysis

Once the archival information was recorded, variance analysis was performed to assess the differences among the links categorized as ‘accessible’, ‘redirected’, ‘broken’ and ‘time out’ as they related to four paper-level metrics: the number of citations in the original paper where the tool was published; number of citations per year in social media platforms such as blogs and Twitter feeds; total readership per year, as measured by Altmetrics; and the Altmetric “attention score.” Because the distributions of all five measures presented heavy tails and deviated from a bell-shaped distribution, we performed a Kruskal–Wallis test on ranks, followed by pairwise Dunn’s tests to confirm which groups presented significant differences with a significance level of 0.01. We provide all p-values and test statistics from these experiments in our electronic supplemental material on GitHub (https://github.com/smangul1/good.software).

## Supplementary Notes

### Supplemental Note 1. An example of the ‘Quick Start’

1. Download the tool using: git clone https://github.com/x/software.tool.git
2. Install tool using: cd software.tool; ./install.sh
3. Run the tool for the example dataset (distributed with the tool): ./software.tool example.dataset

### Supplementary Note 2. Automatic verification of software installability

Software quality, including installability, is typically not thoroughly tested in the formal peer review process, and relying on reviewer feedback can be problematic, as the reviewers may lack the computational skills and time to verify the tools. It is possible to automate the assessment process when software guarantees access to (i) the software binaries or source code; (ii) a script that installs the software in a given Linux environment; (iii) a small example dataset and its expected output; and (iv) a script to perform the analysis on the dataset from *(iii)*.

To provide an automated and openly verifiable certification that a tool is usable, we suggest a model of a server that uses public badges to endorse the installability of a software tool. The server will issue a certificate to the software author, which indicates that the proposed software passed an ‘Automatic Installation Test.’ The installation process, in this case, includes a testing phase that ensures the installation can be successful. Authors of computational tools who submit their software tool to our badge server, alongside an installation script and an example dataset, will receive a badge of confirmation which certifies that the software tool was successfully installed in a third-party environment. Using a Secure Hash Algorithm 45, each generated badge would be unique to each version of the software, installation script, test dataset, and operating system used by the server.

To validate a badge, the server will use a private cryptographic key to publicly sign the badge. Public badge testing provides a strong endorsement of the tool installability up to the current highest standards in the industry, as only the same software version, installation script, and test dataset will confirm the authenticity of the badge and its public signature. A public badge platform will provide a mechanism for researchers and editors of journals in computational biology to verify the installability of a tool in under five minutes through confirmation of the server’s signature. Badges inform the user *a priori* if and under what conditions the software is installable, potentially reducing for each user a significant amount of time that otherwise would be required to test software and attempt installing software that is ultimately uninstallable.

In addition to guaranteeing that a software tool can be successfully installed in a standardized environment, the badge also reflects which specific Linux system was used during the test installation. (Linux-based systems are the most commonly used operating systems in the field of computational biology.) Furthermore, the badge server does not assume an open source software and can be generated based on the source code or binary files.

The server creates an instance of a Linux virtual machine and runs the installation script and test protocol submitted. If the installation completes without errors, and the test dataset provides the expected result, a badge is created that certifies the installability of the tool under the tested conditions. The badge consists of a unique summary generated by the Secure Hash Algorithm 3 (SHA-3) 45, of the items submitted by authors. The server then uses a private cryptographic key to publicly sign this summary. There is a small probability that the hashed summary of two different objects created by SHA-3 could be identical (also known as collision); however, this method is the current technological standard of unique badge creation and is broadly accepted by all industries. Using the server’s public key and the hashed version of the software, any user can authenticate the signature and prove that the server was indeed able to install the tool without manual intervention.

### Supplementary Note 3 List of bioinformatics tools with name Prism

● https://www.ncbi.nlm.nih.gov/pubmed/22851530 (Structural Variance)

● https://academic.oup.com/nar/article/43/20/9645/1394603 (Metabolomics)

● https://www.ncbi.nlm.nih.gov/pubmed/21068001 (Viral Genomics)

● http://honig.c2b2.columbia.edu/prism/ (Protein Structure Analysis) https://www.ncbi.nlm.nih.gov/pubmed/15991339 (Protein Structure)

● https://www.bits.vib.be/software-overview/graphpad-prism (Statistics and visualization)

## Supplementary Tables

**Table S1.** The names of the 10 journals that were used to retrieve the URLs. We reported the total number of papers with URLs in abstract or body of the paper (‘Number of URLs’), and the number of accessible URLs, which were not broken or timeout (‘Number of accessible URLs’).

| Journal name | Number of URLs | Number of accessible URLs |
| --- | --- | --- |
| Nature Biotechnology | 180 | 149 |
| Genome Medicine | 352 | 312 |
| Nature Methods | 403 | 326 |
| Genome Biology | 904 | 746 |
| BMC Systems Biology | 912 | 638 |
| Bioinformatics | 3131 | 2448 |
| PLoS Computational Biology | 3226 | 2529 |
| BMC Bioinformatics | 6840 | 4651 |
| BMC Genomics | 7651 | 5699 |
| Nucleic Acids Research | 13103 | 9011 |

**Table S2.** Installability of 98 published software tools between 2004 and 2018.

| ID | Source | Citations per year | Installation commands executed | Commands documented | Undocumented commands | Automatic installation test | Installation time | Installation difficulty | Sample data provided |
| --- | --- | --- | --- | --- | --- | --- | --- | --- | --- |
| ID1 | Bioconda | 1122.60 | 1 | 1 | 0% | Pass | 5 | Easy | Y |
| ID2 | Bioconda | 13.83 | 1 | 1 | 0% | Pass | 5 | Easy | Y |
| ID3 | Bioconda | 5.14 | 1 | 1 | 0% | Pass | 5 | Easy | Y |
| ID4 | Bioconda | 163.20 | 1 | 1 | 0% | Pass | 5 | Easy | Y |
| ID5 | Bioconda | 247.40 | 1 | 1 | 0% | Pass | 15 | Easy | Y |
| ID6 | Bioconda | 57.20 | 1 | 1 | 0% | Pass | 5 | Easy | Y |
| ID7 | Bioconda | 60.30 | 1 | 1 | 0% | Pass | 5 | Easy | Y |
| ID8 | Other | 33.45 | 3 | 3 | 0% | Pass | 5 | Easy | Y |
| ID9 | Other | 6.83 | 22 | 4 | 82% | Fail | 30 | Complex | Y |
| ID10 | Bioconda | 124.00 | 1 | 1 | 0% | Pass | 30 | Complex | Y |
| ID11 | Other | 1059.00 | 10 | 7 | 30% | Fail | 60 | Complex | Y |
| ID12 | Bioconductor | 159.00 | 20 | 2 | 90% | Fail | 60 | Complex | Y |
| ID13 | pip | 73.17 | 12 | 10 | 17% | Fail | 60 | Complex | Y |
| ID14 | BitBucket | 36.25 | 30 | 10 | 67% | Fail | 120 | Complex | Y |
| ID15 | Bioconductor | 57.83 | 2 | 2 | 0% | Pass | 5 | Easy | Y |
| ID16 | Github | 798.40 | 3 | 2 | 33% | Fail | 5 | Easy | Y |
| ID17 | Bioconductor | 36.00 | 23 | 2 | 91% | Fail | 120 | Complex | Y |
| ID18 | Other | 683.11 | 2 | 2 | 0% | Pass | 5 | Easy | Y |
| ID19 | Github | 216.33 | 12 | 3 | 75% | Fail | 30 | Complex | Y |
| ID20 | Github | 16.25 | 8 | 6 | 25% | Fail | 60 | Complex | Y |
| ID21 | Other | 14.70 | 13 | 13 | 0% | Pass | 15 | Easy | Y |
| ID22 | SourceForge | 22.71 | 190 | 7 | 96% | Fail | 120 | Not installed | Y |
| ID24 | Other | 169.22 | 2 | 2 | 0% | Pass | 5 | Easy | Y |
| ID25 | Other | 15.63 | 52 | N/A | 0% | Fail | 120 | Complex | N |
| ID26 | BitBucket | 0.71 | 7 | 4 | 43% | Fail | 30 | Complex | Y |
| ID27 | Other | 39.10 | 12 | 12 | 0% | Pass | 15 | Easy | Y |
| ID28 | Other | 12.75 | 4 | 4 | 0% | Pass | 15 | Easy | Y |
| ID29 | Other | 1.71 | 4 | 4 | 0% | Fail | 30 | Not installed | Y |
| ID30 | Other | 1.00 | 2 | 2 | 0% | Pass | 15 | Easy | Y |
| ID31 | Other | 0.29 | 7 | 2 | 71% | Fail | 5 | Not installed | Y |
| ID32 | Other | 2.60 | 11 | 4 | 64% | Fail | 15 | Not installed | Y |
| ID33 | Other | 13.33 | 5 | 5 | 0% | Pass | 15 | Easy | Y |
| ID34 | Other | 3.20 | 5 | 3 | 40% | Fail | 30 | Complex | Y |
| ID35 | Other | 8.00 | 4 | 3 | 25% | Fail | 15 | Easy | Y |
| ID36 | Other | 2.50 | 7 | 7 | 0% | Fail | 15 | Not installed | Y |
| ID37 | Other | 3.00 | 6 | 6 | 0% | Pass | 15 | Easy | Y |
| ID38 | Other | 3.25 | 5 | 4 | 20% | Fail | 15 | Easy | Y |
| ID39 | Other | 3.00 | 10 | 10 | 0% | Fail | 30 | Not installed | Y |
| ID40 | Other | 2.67 | 3 | 3 | 0% | Pass | 15 | Easy | Y |
| ID41 | Other | 1.33 | 2 | 2 | 0% | Pass | 5 | Easy | Y |
| ID42 | Other | 124.10 | 1 | 1 | 0% | Pass | 5 | Easy | N |
| ID43 | Other | 15.30 | 1 | 1 | 0% | Pass | 5 | Easy | Y |
| ID44 | Other | 7.17 | 2 | 1 | 50% | Fail | 15 | Easy | N |
| ID45 | SourceForge | 1.83 | 1 | N/A | 0% | Fail | 15 | Not installed | Y |
| ID46 | Github | 1.50 | 1 | 1 | 0% | Pass | 15 | Easy | N |
| ID47 | Github | 1.33 | 3 | 3 | 0% | Pass | 5 | Easy | Y |
| ID48 | Github | 2.75 | 7 | 7 | 0% | Pass | 15 | Easy | Y |
| ID49 | Github | 3.00 | 7 | 7 | 0% | Pass | 15 | Easy | N |
| ID50 | Bioconductor | 1.00 | 1 | 1 | 0% | Pass | 5 | Easy | Y |
| ID51 | Github | 0.67 | 5 | 3 | 40% | Fail | 30 | Not installed | Y |
| ID52 | Other | 39.00 | N/A | N/A | 0% | Pass | 5 | Easy | Y |
| ID53 | Bioconductor | 26.57 | 2 | 1 | 50% | Fail | 5 | Easy | N |
| ID54 | Other | 43.00 | 5 | 1 | 80% | Fail | 120 | Complex | Y |
| ID55 | SourceForge | 8.00 | 1 | 1 | 0% | Pass | 5 | Easy | N |
| ID56 | SourceForge | 16.00 | 2 | N/A | 0% | Fail | 5 | Easy | N |
| ID57 | Other | 4.67 | 1 | 1 | 0% | Pass | 5 | Easy | N |
| ID58 | Github | 13.75 | 5 | N/A | 0% | Fail | 30 | Complex | N |
| ID59 | Other | 2.43 | 2 | N/A | 0% | Fail | 5 | Easy | N |
| ID60 | SourceForge | 1.60 | 5 | N/A | 0% | Fail | 30 | Complex | N |
| ID61 | Github | 1.67 | 1 | 1 | 0% | Pass | 5 | Easy | Y |
| ID62 | Github | 5.75 | 4 | 3 | 25% | Fail | 120 | Complex | N |
| ID63 | Github | 1450.90 | 3 | 3 | 0% | Pass | 5 | Easy | Y |
| ID64 | Other | 30.75 | 7 | 3 | 57% | Fail | 120 | Complex | Y |
| ID65 | Github | 3.75 | 3 | 2 | 33% | Fail | 5 | Easy | N |
| ID66 | Other | 1.33 | 1 | 1 | 0% | Pass | 5 | Easy | N |
| ID67 | Other | 24.00 | 4 | N/A | 0% | Fail | 120 | Not installed | N |
| ID68 | Other | 15.10 | 4 | 2 | 50% | Fail | 120 | Complex | Y |
| ID69 | Other | 4.33 | 4 | N/A | 0% | Fail | 120 | Not installed | N |
| ID70 | SourceForge | 13.29 | 1 | 1 | 0% | Fail | 120 | Not installed | N |
| ID71 | Other | 16.29 | 1 | 1 | 0% | Pass | 5 | Easy | Y |
| ID72 | Other | 9.56 | N/A | N/A | 0% | Fail | 120 | Not installed | Y |
| ID73 | Other | 7.25 | N/A | 3 | 0% | Fail | 120 | Not installed | N |
| ID74 | Other | 5.11 | N/A | N/A | 0% | Fail | 120 | Not installed | N |
| ID75 | Other | 9.80 | N/A | 1 | 0% | Fail | 120 | Not installed | Y |
| ID76 | Other | 7.00 | N/A | N/A | 0% | Fail | 120 | Not installed | N |
| ID77 | Github | 4.50 | N/A | 6 | 0% | Fail | 120 | Not installed | N |
| ID78 | SourceForge | 3.00 | 9 | 9 | 0% | Fail | 120 | Not installed | Y |
| ID79 | Other | 47.13 | N/A | 2 | 0% | Fail | 5 | Not installed | N |
| ID80 | Other | 1.38 | N/A | N/A | 0% | Fail | 5 | Not installed | N |
| ID81 | Other | 3.33 | N/A | N/A | 0% | Fail | 120 | Not installed | Y |
| ID82 | Github | 3.33 | 6 | 6 | 0% | Fail | 120 | Not installed | Y |
| ID83 | Other | 8.90 | 14 | 14 | 0% | Fail | 120 | Not installed | Y |
| ID84 | Other | 23.60 | N/A | N/A | 0% | Fail | 30 | Not installed | Y |
| ID85 | Other | 24.20 | N/A | 7 | 0% | Fail | 120 | Not installed | Y |
| ID86 | Other | 28.50 | N/A | 29 | 0% | Fail | 120 | Not | N |
|  |  |  |  |  |  |  |  | installed |  |
| ID87 | SourceForge | 10.56 | N/A | N/A | 0% | Pass | 5 | Easy | N |
| ID88 | SourceForge | 12.25 | N/A | N/A | 0% | Pass | 5 | Easy | N |
| ID89 | Other | 10.43 | N/A | N/A | 0% | Pass | 120 | Complex | Y |
| ID90 | Other | 12.17 | 1 | 1 | 0% | Pass | 5 | Easy | Y |
| ID91 | Other | 2.00 | 2 | 2 | 0% | Pass | 5 | Easy | N |
| ID92 | Github | 2.25 | 12 | 12 | 0% | Pass | 120 | Complex | Y |
| ID93 | Other | 1.67 | 3 | 3 | 0% | Pass | 5 | Easy | N |
| ID94 | Github | 3.25 | 1 | 1 | 0% | Pass | 5 | Easy | Y |
| ID95 | Github | 16.89 | 10 | 3 | 70% | Fail | 120 | Not installed | Y |
| ID96 | BitBucket | 9.71 | 3 | 3 | 0% | Pass | 120 | Complex | N |
| ID97 | SourceForge | 76.30 | 1 | 1 | 0% | Pass | 5 | Easy | Y |
| ID98 | Github | 3.00 | 9 | 9 | 0% | Pass | 5 | Easy | Y |
| ID99 | Github | 75.71 | 1 | 1 | 0% | Pass | 5 | Easy | N |

**Table S3.**
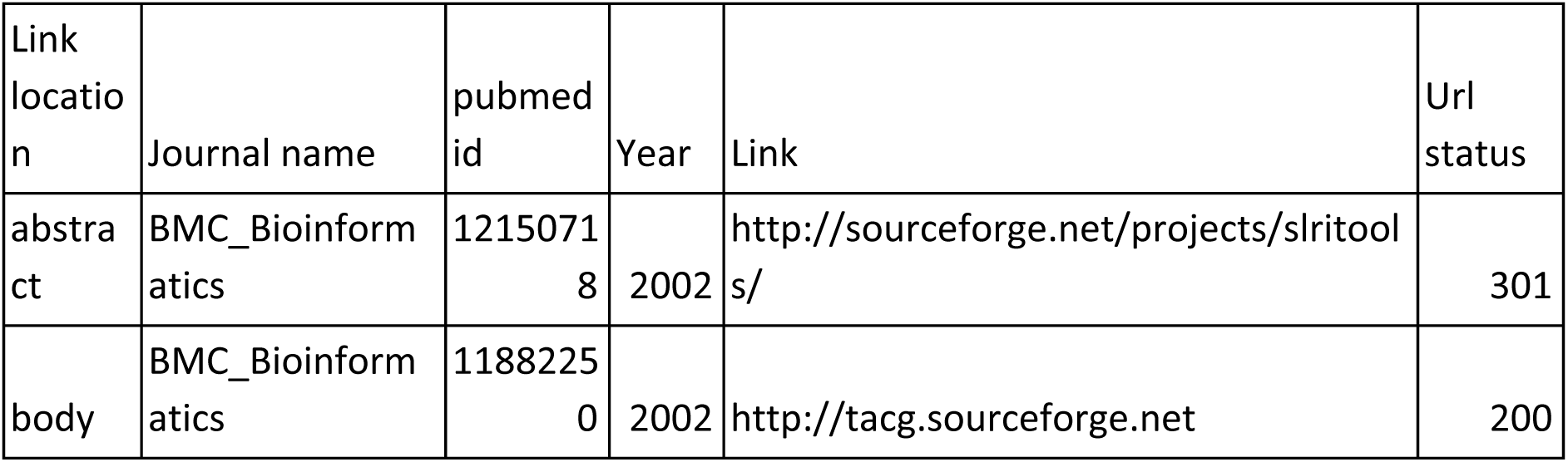
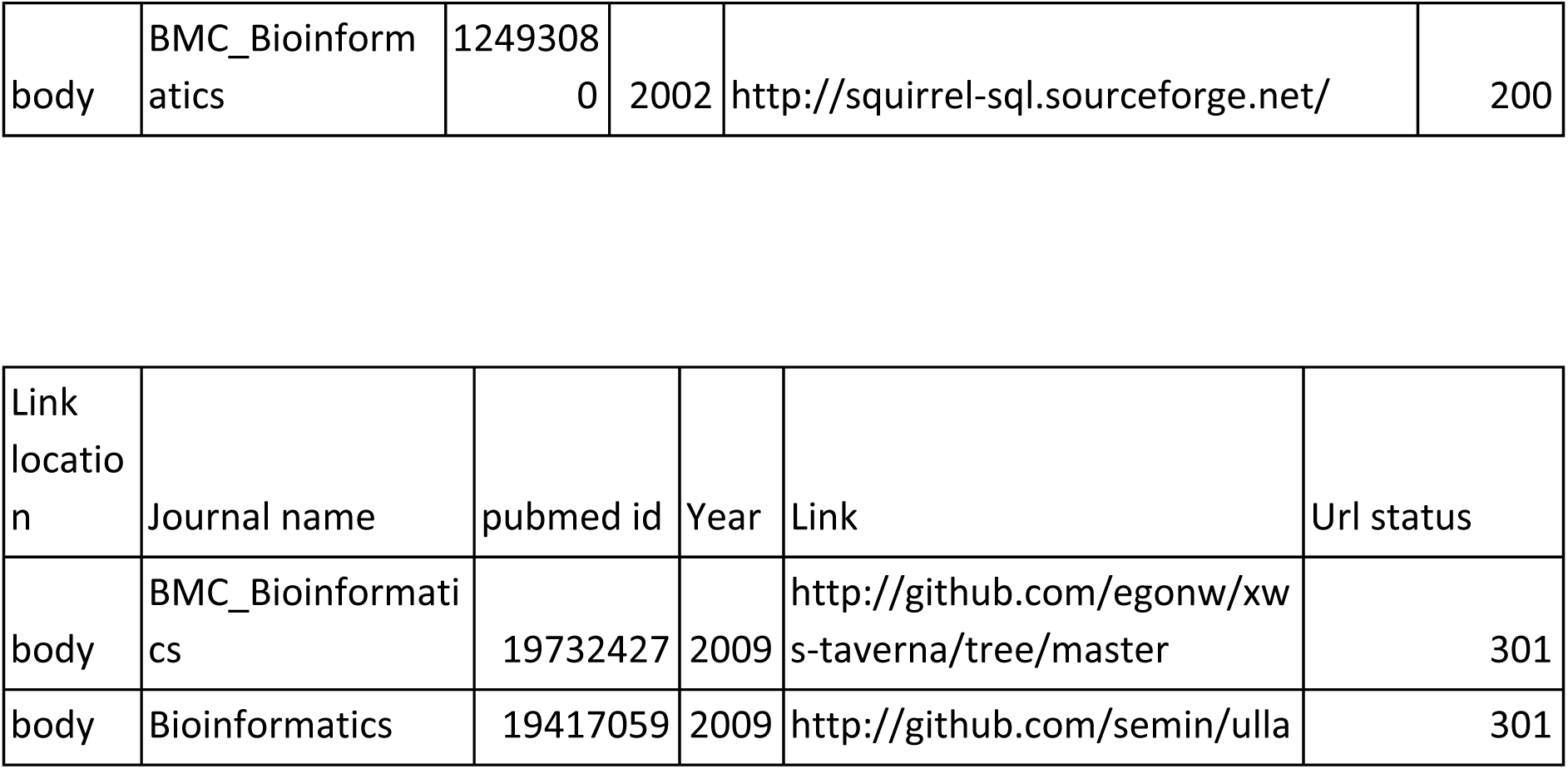
List of earliest published software tools and resources stored on https://sourceforge.net and https://github.com/

## Supplementary Figures

**Figure S1.**
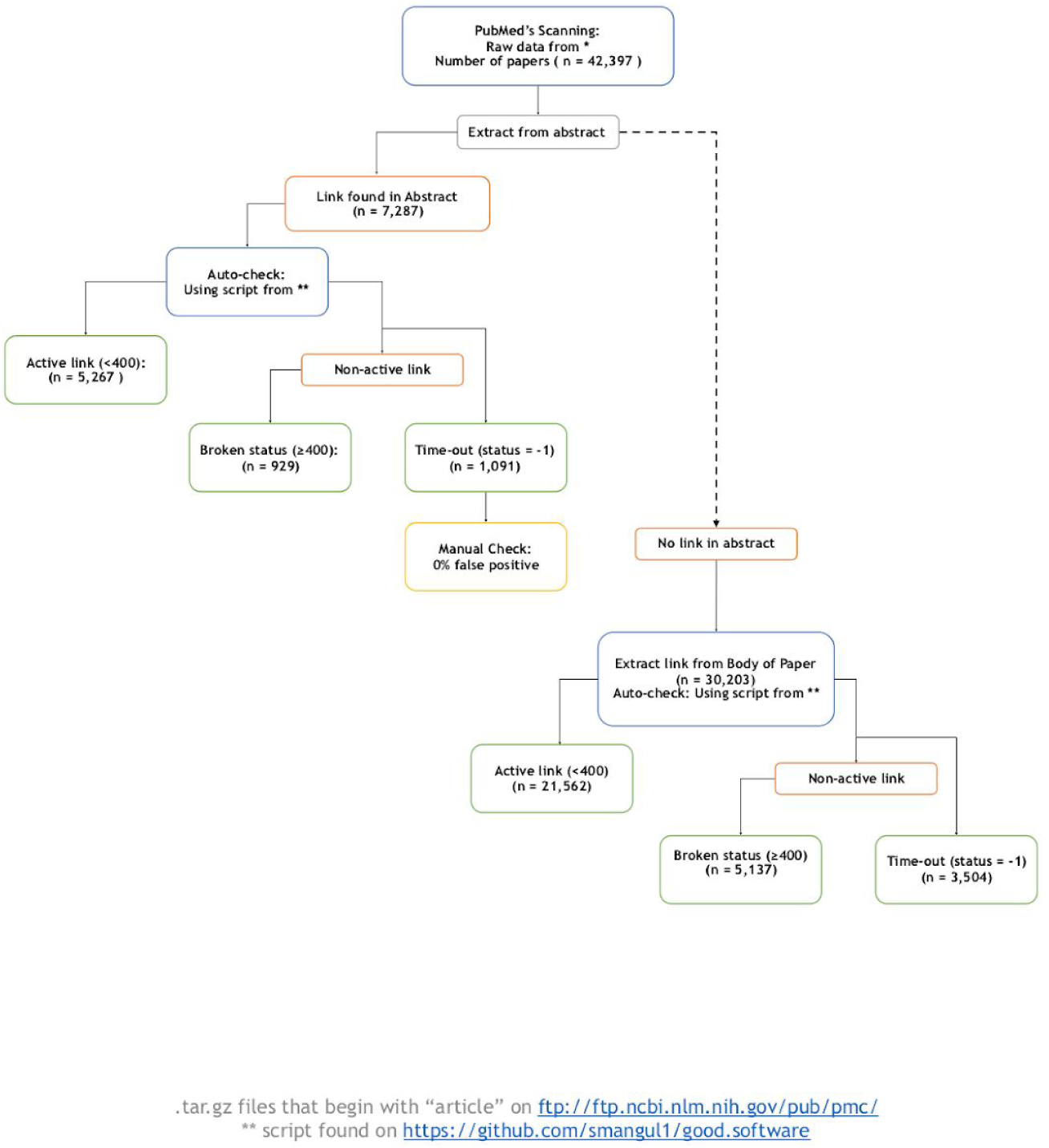
Protocol to check the archival stability of a published software tool or resource. Numbers are provided for illustrative purposes and correspond to the link presented in the abstracts of the published papers considered in this study.

**Figure S2.**
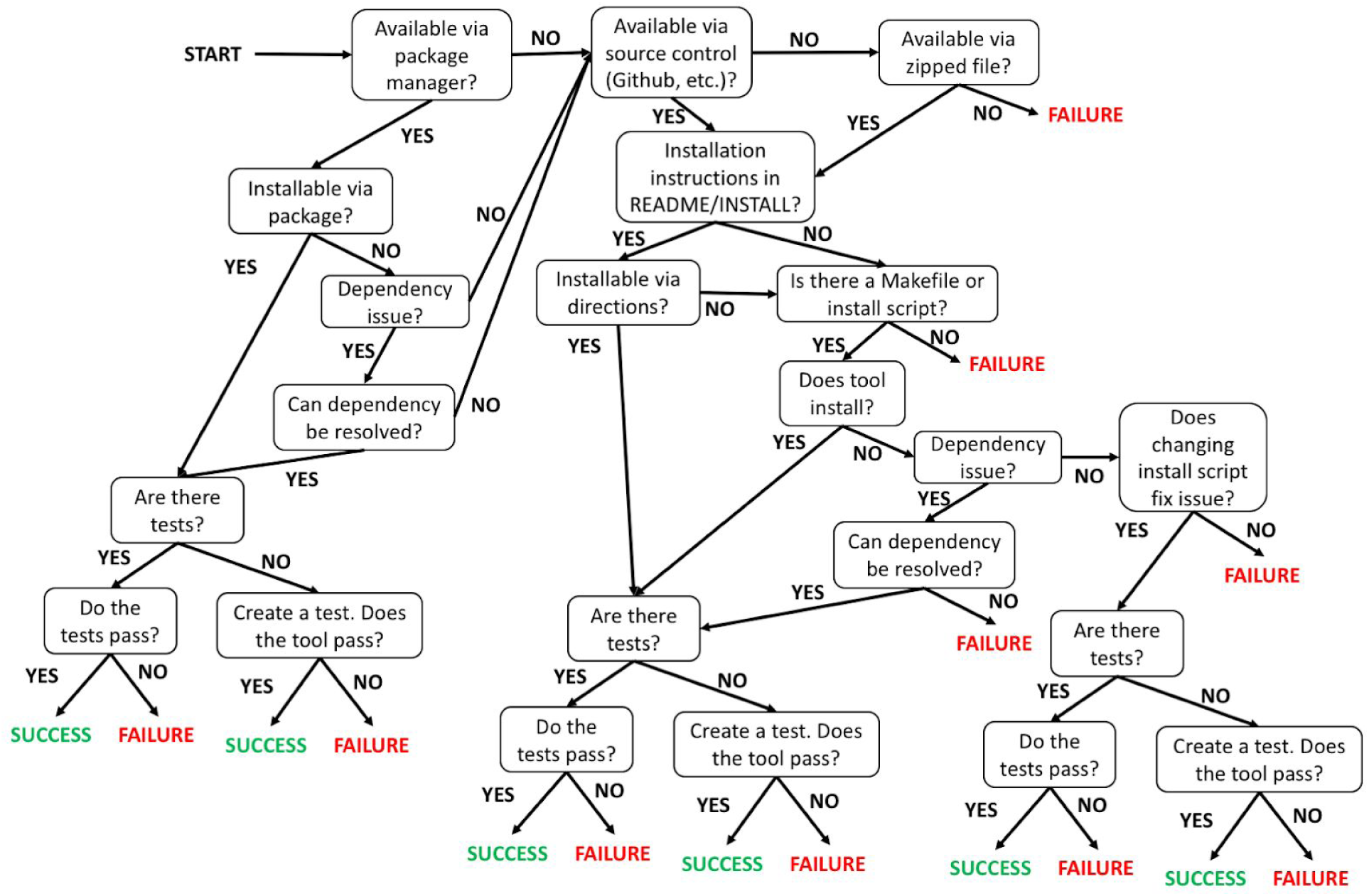
Protocol to verify the installability of a published software tool.

**Figure S3.**
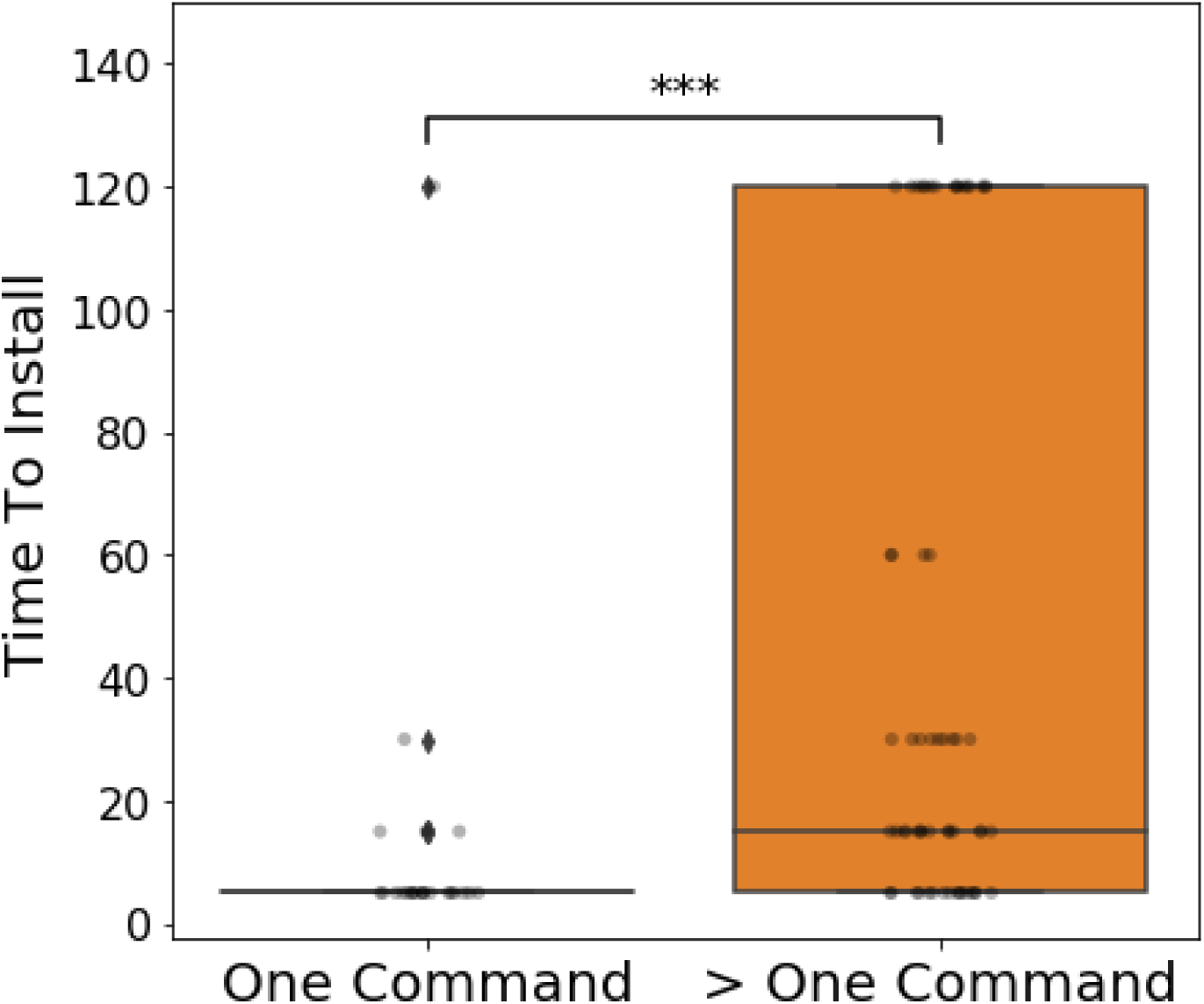
A box plot showing the time required to install tools that required a single command, compared to tools that required multiple (Mann–Whitney *U* test, p-value=4.7×10^−6^).

